# Sniffing Out New Friends: Similarity in Body-Odor Predicts the Quality of Same-Sex Non-Romantic Dyadic Interactions

**DOI:** 10.1101/2021.06.14.448352

**Authors:** Inbal Ravreby, Kobi Snitz, Noam Sobel

## Abstract

Most are familiar with the notion of socially “clicking” with someone, namely sensing an immediate bond that can lead to strong and often long-lasting friendships. The mechanisms underlying such rapid bonding remain unclear. Given that body-odor similarity is a critical cue for social interaction in non-human mammals, we tested the hypothesis that body-odor similarly contributes to bonding in same-sex non-romantic human dyads. We observed that objective ratings obtained with an electronic nose, and subjective ratings obtained from human smellers, converged to suggest that click-friends smell more similar to each other than random dyads. Remarkably, we then found that we could use the electronic nose to predict which strangers would later form better dyadic interactions. Thus, humans may literally sniff-out new friends based on similarities in body-odor.

## Introduction

Human dyadic same-sex non-romantic friendships are a critical pillar of psychological health (*1*, *2*). Such friendships can develop slowly over time, but occasionally, in so-called click-friendships, a strong sense of bonding can form almost instantaneously (*3*). Because similarity within a dyad is a strong positive predictor of friendship (*4*–*8*), that acts at the very early phase of interaction (7), one may assume that similarity also plays a role in forming such click-friendships. The known similarities that predict friendship range from the unsurprising such as age, race, education, religion, and indeed, physical appearance (*9*, *10*), on to more complex measures such as personality (*11*, *12*) and values (*13*), and culminating in measures such as patterns of neural activity (*14*), and genetic makeup (*15*–*18*). Non-human mammals rapidly obtain complex social information from body-odor (*19*). Humans also constantly sniff conspecifics (*20*), and themselves (*21*), to obtain social information, and similarity in the sniffed body-odor infers kinship with self (*22*, *23*), or between strangers (*22*, *24*). Moreover, growing evidence implies that humans can infer the emotional state of conspecifics, whether it be fear (*25*) or happiness (*26*), from body-odor alone. Given that a friend’s body-odor and one’s own body-odor induce similar patterns of brain activity, yet exposure to a stranger’s body-odor induces a very different limbic fear-type brain response (*27*), we hypothesized that similarity in body-odor may contribute to rapid friendship formation. To test this, we first asked whether click-friends indeed smell alike. After recruiting same-sex non-romantic click-friends and harvesting their body-odor, we found that both an analytical device (an electronic nose) and independent human smellers, converged to rate the body-odors of click-friends as more similar than those of random dyads. Finally, to ask whether similarity in body-odor is not merely a consequence of friendship, we used the electronic nose to predict social interactions between strangers. We found that strangers whose body-odor was more similar were more prone to later positive dyadic social interaction. Thus, we conclude that similarity in human body-odor is related to a fundamental mechanism involved in friendship formation.

## Results

### Defining click-friendships

Although clicking is an intuitively clear term in the context of friendship (*3*), we are unaware of a formal definition for it in the literature. To define click-friendship, in Experiment 1, 235 participants (135 women, aged between 20 and 43 years, M = 26.35 ± 4.166) were asked to define what click-friendship is in their own words. Only 10 of 235 participants said they did not know what click-friendship is. That 225 of 235 participants had a clear notion of what we were asking about, further supports that click-friendship is a real social event, despite the lack of formal definition. Moreover, we observed high consistency across individuals, whereby the 235 participants spontaneously converged to use only 42 meaningfully different statements to define click-friendship. These statements and their frequency of application are in Table S1, and we used the top 20 descriptors as our conditioned definition of click-friendships.

### The body-odors of click-friends are more similar than expected by chance

We conducted a 6-month long social-media-centered recruitment effort in search of click-friends who then retained a lasting relationship. After phone interviews and questionnaires, this culminated in 20 same-sex non-romantic click-friend dyads (10 male, aged between 22 and 39 years, M ± SD = 24.757 ± 3.388, mean friendship duration = 6.185 ± 5.793 years) from all over Israel. These dyads mutually self-reported that their friendship began as a click-friendship, and each affirmed all 20 top criteria of Table S1. These participants donated body-odor using a strict body-odor-donation protocol (see Methods). To ask whether there is similarity in the body-odor chemical fingerprint across members of click-dyads, in Experiment 2 we first sampled all body-odors with an electronic nose (eNose) (Fig. 1A). This particular eNose (PEN3, AirSense Analytics, Schwerin, Germany) has 10 metal oxide sensors, each coated with a different material conferring chemical specificity. Thus, each sample is potentially made of 10 responses that combine to generate a specific pattern associated with an odor. We observed that only 5 of the sensor subtypes responded to body-odors (Sensors # 2, 6, 7, 8 and 10). We thus represented each of the 40 body-odors as a five-dimensional vector (all Experiment 2 raw eNose results available in Data File 1). We then calculated the Euclidian distance between the two body-odors from each dyad within the five-dimensional space. We observed that mean Euclidian distance between click-friends was 5.075 ± 4.947 arbitrary units (AU) within the five-dimensional space. In turn, we used the same 40 individuals to randomly generate 10,000 iterations of 20 same-sex dyads to obtain a distribution of the mean Euclidean distances between 20 random dyads. We observed that mean Euclidian distance between such random same-sex dyads was 6.535 ± 0.55 AU. Using a bootstrap test, we find that these values are significantly different (mean clicks = 5.075 ± 4.947, mean random dyads = 6.535 ± 0.55, bootstrapped p = 0.0059, Cohen’s d of bootstrap statistic = 2.654), or in other words, the chemical signature from body-odors of click-friends is significantly more similar than the chemical signature from body-odors of random dyads (Fig. 1B).

**Fig. 1.**
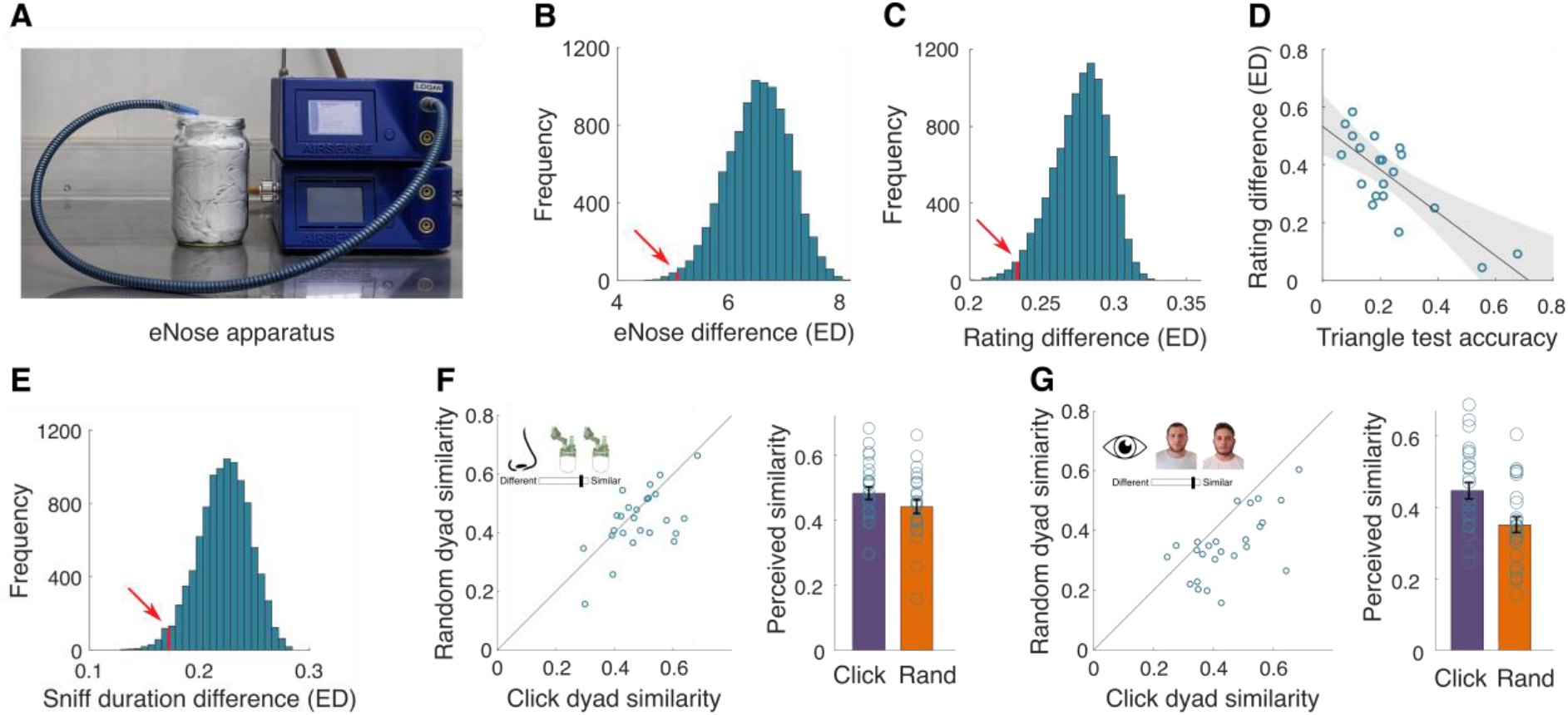
The body-odors of click-friends are more similar than expected by chance. (**A**) A PEN3 eNose was used to measure headspace over a T-shirt in a jar. (**B**) Histogram showing 10,000 iterations of the average Euclidian distance (ED) between 20 same-sex random dyads in the eNose-space. The distance between click-friends is denoted by the red line and arrow. (**C**) Histogram showing 10,000 iterations of the average Euclidian distance (ED) between 20 same-sex random dyads in perceptual rating space. The distance between click-friends is denoted by the red line and arrow. (**D**) Pearson correlation between the difference in perceptual ratings and triangle test accuracy (n = 20). The black line is the linear regression, and the grey area marks the CI of the regression line. (**E**) Histogram showing 10,000 iterations of the average Euclidian distance (ED) between 20 same-sex random dyads in sniff duration. The distance between click-friends is denoted by the red line and arrow. (**F**) Perceived perceptual odor similarity for click-dyads (X axis) vs. random dyads (Y axis). Each point is a rater (n = 25), and the point reflects the average of their 40 ratings (20 click, 20 random). The diagonal line reflects the unit slope line (x = y), such that if points accumulate under the line then the values are greater for click dyad similarity, and if they accumulate above the line then the values are greater for random dyad similarity. The associated bar graph is the average perceived similarity. (**G**) This panel is identical to panel F, but for visual rather than olfactory similarity data.

Chemical similarity inferred by eNose does not necessarily imply human perceived olfactory similarity. To ask whether the eNose results are mirrored in human perception, in Experiment 3 we recruited 24 smellers (13 females, aged between 22 and 39 years, M ± SD = 27 ± 4.63). We designed the perceptual experiment such that it would tap both explicit and implicit classification. To probe explicit classification, we used a triangle test. On each trial, the participant was presented with a body-odor triplet, where two odorants were from members of a click dyad, and the third distractor odorant was from an unrelated same-sex body-odor donor. Participants were asked to select the odorant outlier. Each participant completed 20 trials (inter-trial-interval = 25 s), one for each click dyad. In an ensuing task, participants performed the very same classification using face-torso photographs rather than body-odors. Next, to probe implicit classification, participants smelled the 40 click-friend body-odors one-by-one, randomly ordered, and rated them using visual analog scales (VASs) for pleasantness, intensity, sexual attraction, competence, and warmth (temperament). Whereas the former two descriptors were used because they reflect the primary dimensions of odor (*28*), the latter two descriptors were used because they reflect primary dimensions in social interaction (*29*). Finally, given that sniffing patterns provide an added implicit measure of olfactory perception (*30*), throughout all tasks participants wore a nasal cannula linked to a spirometer, providing a precise measure of nasal airflow (all Experiment 3 raw data is available in Data File 2).

Significant classification is typically attributed to a d-prime (d’) score of “1” or higher (*31*, *32*), and in the triangle test where chance = 33.33%, d’= 1 is at 41.8% accuracy^19,20^. We observe that overall for the group of click-dyads, mean accuracy was 36.15% ± 14.54 (t(19) = 0.894, p = 0.382, Cohen’s d = 0.212), reflecting a mean d’ score of 0.693 ± 0.69 (one sample two tailed t-test against d’ = 1: t(19) = 1.99, p = 0.061, Cohen’s d = 0.445). In other words, at the group-level, participants failed to significantly classify click-dyads based on body-odor. Similarly, albeit a trend, participants failed to significantly classify click-dyads based on photographs in a triangle test (mean accuracy = 42.29% ± 20.6%, t(19) = 2.017, p = 0.058, Cohen’s d = 0.445, reflecting a mean d’ score of 1.131 ± 0.979, t(19) = 0.598, p = 0.57, Cohen’s d = 0.134).

To examine implicit perceived body-odor similarity, we compared the similarity within click-dyads vs random dyads in the five-dimensional VAS space. This was done by comparing the average distance between the 20 click-dyads to the probability distribution of the average distance between 20 same-sex random dyads, 10,000 times. We found that the Euclidian distances were significantly lower between click-dyads compared to random dyads (mean clicks = 0.233 ± 0.153, mean random = 0.277 ± 0.019, bootstrapped p = 0.0187, Cohen’s d of bootstrap statistic = 2.316) (see Fig. 1C). To ask whether this effect was carried by any particular descriptor, we repeated the analysis for each descriptor alone. We found that click-dyads were rated as significantly more similar in body-odor pleasantness (mean clicking = 0.1 ± 0.11 AU, mean random = 0.128 ± 0.013 AU, bootstrapped p = 0.03, Cohen’s d = 2.154), in body-odor attractiveness (mean clicking = 0.096 ± 0.106 AU, mean random = 0.13 ± 0.013 AU, bootstrapped p = 0.01, Cohen’s d = 2.615) and in the competence associated with the body-odor (mean clicking = 0.065 ± 0.063 AU, mean random = 0.084 ± 0.009 AU, bootstrapped p = 0.027, Cohen’s d = 2.11) than random dyads (see Fig. S1 in the Supplementary Materials). Moreover, despite no overall group-effect in the previous triangle test, we observed that the better a given click-dyad was explicitly classified in the triangle test, the more similar their body-odors were in the implicit rating experiment (Pearson r = −0.785, p = 0.000041) (Fig. 1D). Finally, given that sniff duration is modulated in accordance with odorant content (*33*), we compared sniff duration across samples. We observed significantly greater similarity in sniff duration when sniffing members of a click-dyad versus sniffing random dyads (mean clicks’ duration difference = 0.171 ± 0.173 seconds, mean random = 0.222 ± 0.023, bootstrapped p = 0.0237, Cohen’s d = 2.217) (Fig. 1E).

Whereas the above implicit measures suggested that click-friends indeed smell alike, the previous explicit triangle test did not. The triangle test, however, entails an inherent memory component that may complicate the comparison of body-odors. To address this, in Experiment 4 we conducted a different explicit test, where now 25 participants (19 females, aged between 21 to 38 years, M = 25.76, ± 4.075) explicitly rated the perceptual similarity of pairs of body-odors. Each participant rated the perceptual similarity of 40 dyads, 20 click-dyads, and 20 random same-sex dyads, along a VAS ranging from *similar* to *different*. An additional 25 participants (13 females, aged between 21 to 37 years, M = 26.56 ± 5.091) conducted a similar experiment using face-torso photographs rather than body-odors (All Experiment 4 raw data is available in data file 3). We observed that the body-odors of click-friends were significantly more explicitly similar to each other than the body-odors of random dyads (mean clicks = 0.483 ± 0.0972, mean random = 0.442 ± 0.105, two-tailed paired t-test: t(24) = 2.206, p = 0.0372, Cohen’s d = 0.4). Consistent with previous reports (*10*), we also observed remarkably greater visual similarity between click-friends using this paradigm (mean click = 0.447 ± 0.116 VAS units, mean random = 0.351 ± 0.11 VAS units, two-tailed paired t-test: t(24) = 4.7967, p = 6.9633e-05, Cohen’s d = 0.84) (Fig. 1F-G). In other words, unlike the triangle test, this explicit test converged with the implicit measures to suggest that click-dyads smell alike, well beyond random dyads.

Both eNose similarity and human perceptual similarity were higher for click-friends vs. random dyads. This outcome may imply that either the eNose provides a good reflection of human perception, or in turn that each measurement type, eNose and perception, captured a different portion of the chemical variance underlying this link. To address these alternatives, we asked whether eNose-derived similarity was correlated with perceptual similarity across the 20 dyads. We observed no sign of such correlation (Pearson r = −0.16, p = 0.51, and after removing an outlier, r = −0.29, p = 0.23) (Fig. S2), implying that the chemical cues used by the eNose were likely not those used by human raters.

### An electronic nose can predict social interaction

The above results suggest that click-friends have greater similarity in body-odor chemistry and in body-odor perceived smell in comparison to random dyads. This similarity may somehow be a consequence of friendship (common body-odor-shaping experiences, e.g., living in the same area, eating together), or it may be related to the root causes of friendship. To disentangle these alternatives, in Experiment 5 we tested whether similarity in body-odor as determined by eNose can predict the quality of social interaction between complete strangers. We recruited 17 strangers (10 females, ages between 20 and 37 years, M = 26.2 ± 4.56), and collected their body-odors as before. To force non-verbal dyadic interaction, we used the Mirror Game (*34*). In this paradigm, two participants stand facing each other 50 cm apart (i.e., a close distance allowing body-odor exposure), and for 2 minutes try to mirror each-other’s hand motion (Fig. 2A). Participants were not allowed to talk throughout the experiment. This paradigm provides for several potential measures on the quality of dyadic interaction. First, using motion energy analysis (*35*), we can calculate the extent or accuracy of mirroring. Second, after each 2-minute interaction, participants rated their partner using a continuous version of the “Inclusion of Other in the Self” (IOS) scale (*36*), where participants place two circles to graphically represent the quality of “overlap” with their partner. Third, the participants rated the quality of interaction with their partner along 12 relevant VASs (Fig. 3). Fourth, participants were requested to indicate whether they clicked with their partner or not (this binary indication is not taken to imply that they here became click-friends). We applied a within-sex round-robin design such that each participant played with each of the other same-sex participants, providing for 66 novel dyads, 21 male, and 45 female (All Experiment 5 raw data is available in Data File 4). eNose analyses were performed after completion of the experiment, rendering all interactions double-blind. We calculated as before the eNose-derived chemical similarity between all 66 possible same-sex novel dyads (all raw eNose results available in Data File 4). We observe that 22 dyads (8 male and 14 female) reported a mutual click. We compared the eNose distance between these 22 dyads to the eNose distance between 10,000 random selections of 22 (out of the 66) same-sex non-mutual-clicking dyads, and observed that dyads who reported clicking were significantly more chemically similar than dyads that did not report clicking (mean clicking = 1.592 ± 0.803 AU, mean random = 2.003 ± 0.1559 AU, bootstrapped p = 0.0029, Cohen’s d of bootstrap statistic = 2.636) (Fig. 2B, 2C, 2D).

**Fig. 2.**
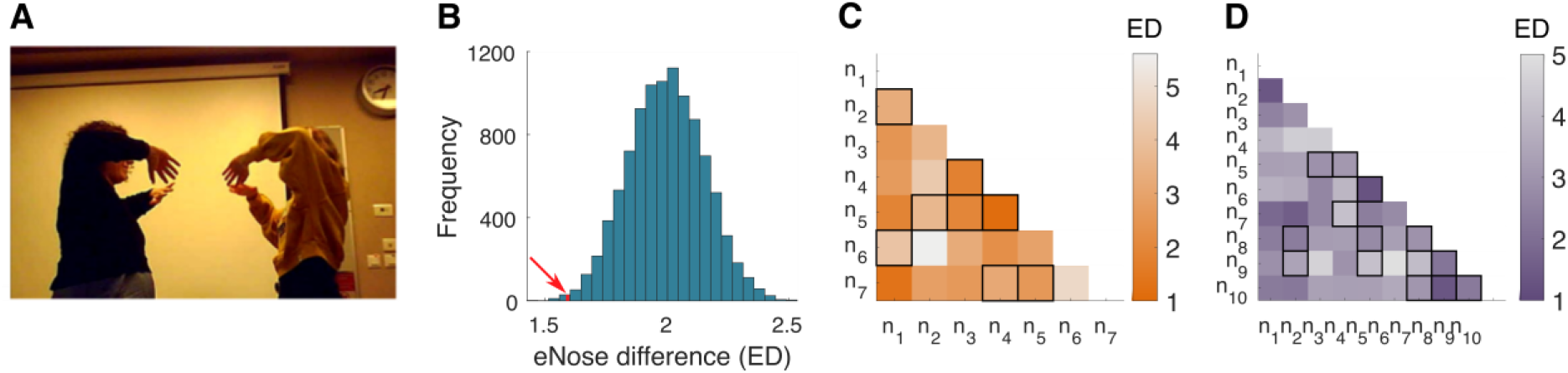
Body-odor similarity is related to clicking in the Mirror Game. (**A**) A dyad playing the mirror game. (**B**) Histogram showing 10,000 iterations of the average Euclidian distance (ED) between 22 same-sex random dyads who played the mirror game, represented in the eNose-space. The distance between the 22 dyads who reported clicking in the game is denoted by the red line and arrow. (**C**) eNose-derived Euclidean distance (ED) between all male dyads (n = 21) who played the mirror game. Dyads who reported a mutual click are outlined in black. (**D**) eNose-derived Euclidean distance (ED) between all female dyads (n = 45) who played the mirror game. Dyads who reported a mutual click are outlined in black.

**Fig. 3.**
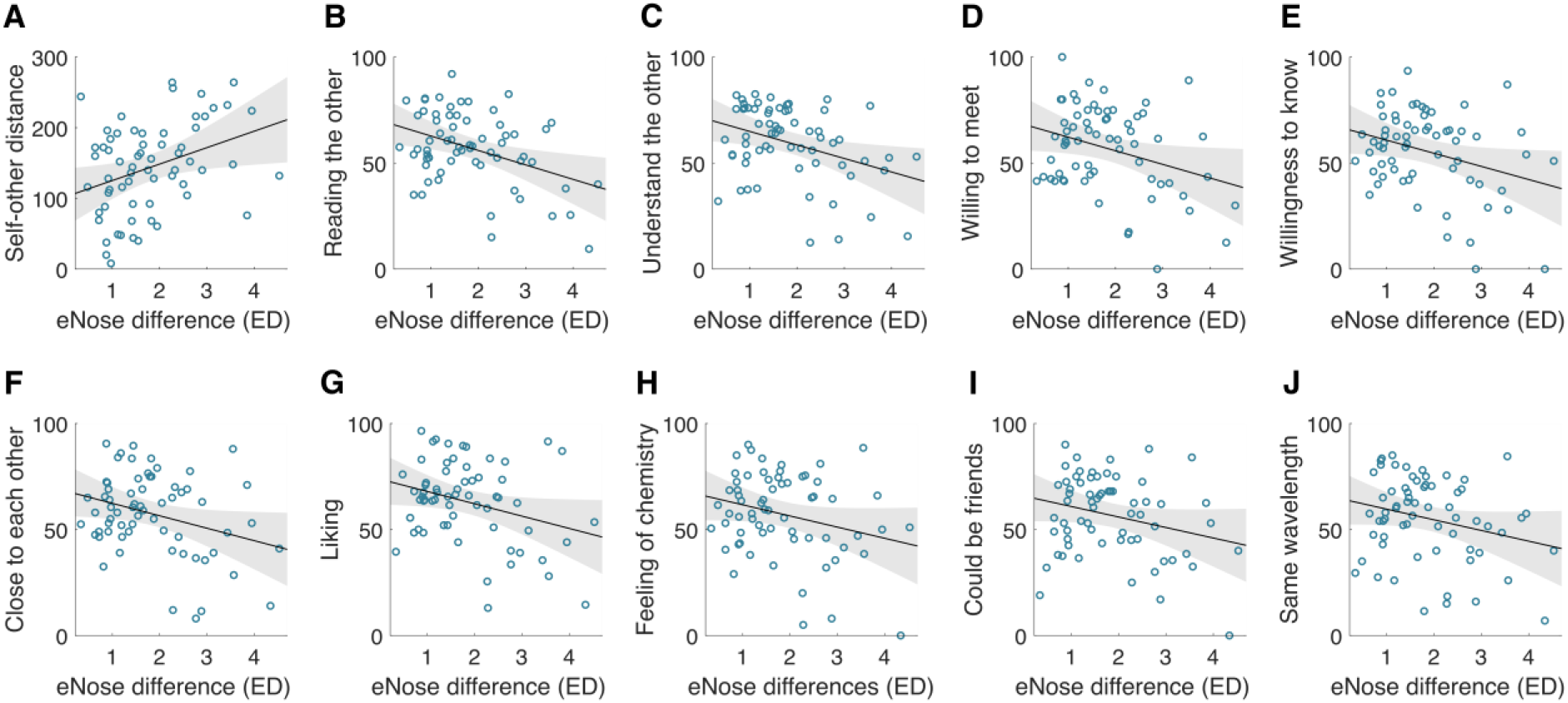
Body-odor similarity is related to the quality of interaction in strangers. Each panel is the Pearson correlation between the chemical difference in a dyad as determined by eNose vs. one of the 13 measures of social interaction. Each blue circle is one of the 66 that played the mirror game. The black line is the linear regression line and the grey area marks the CI of the regression. The 13 measures are: (A) including the other in the self as was measured in the IOS, (B) reading the partner’s mind, (C) understanding the partner, (D) willingness to meet again with the partner, (E) willingness to get to know the partner, (F) feeling close to the partner, (G) liking the partner, (H) feeling chemistry with the partner, (I) thinking that they could be good friends, (J) feeling on the same wavelength, (K) feeling as if they already knew the partner, (L) feeling comfortable to share personal issues with the partner and (M) feeling that the partner was friendly.

Moreover, we observed that the eNose-derived chemical similarity between dyad members was significantly correlated (FDR corrected) with IOS scores, as well as with 9 of the remaining 12 measures of interaction provided by participants (correlation between eNose-derived chemical similarity (ED) and IOS score: r = 0.35, p = 0.0046, Benjamini-Hochberg P = 0.02. VAS scales: “reading the partner’s mind”: r = −0.4, p = 0.001, Benjamini-Hochberg P = 0.013; “understanding the partner”: r = −0.35, p = 0.0028, Benjamini-Hochberg P = 0.018; “willingness to meet again with the partner”: r = −0.32, p = 0.0095, Benjamini-Hochberg P = 0.025; “willingness to get to know the partner”: r = −0.32, p = 0.011, Benjamini-Hochberg P = 0.025; “feeling close to the partner”: r = −0.31, p = 0.012, Benjamini-Hochberg P = 0.025; “liking the partner”: r = −0.31, p = 0.014, Benjamini-Hochberg P = 0.025; “feeling chemistry with the partner”: r = −0.27, p = 0.032, Benjamini-Hochberg P = 0.047; “thinking that they could be good friends”: r = −0.26, p = 0.033, Benjamini-Hochberg P = 0.047; “feeling on the same wavelength”: r = −0.26, p = 0.036, Benjamini-Hochberg P = 0.047; “feeling as if they already knew the partner”: r = −0.205, p = 0.099, Benjamini-Hochberg P = 0.116; “feeling comfortable to share personal issues with the partner”: r = −0.2, p = 0.11, Benjamini-Hochberg P = 0.122; and “feeling that the partner was friendly”: r = −0.15, p = 0.25, Benjamini-Hochberg P = 0.25. (Fig. 3).

In turn, we did not observe any link between the quality of mirroring as estimated by motion energy analysis and eNose-derived chemical similarity. This null result was likely influenced in part by a sex-difference whereby female-dyads achieved high mimicking scores yet male-dyads achieved low mimicking scores (Fig. S2). In combination, the above results imply that the more chemically similar the dyad body-odors were, the better they interacted. Given these relationships, we asked whether a classifier could use eNose data to predict social interaction. Using a weighted KNN classifier, we contrasted the 22 dyads that mutually reported clicking, with the other 44 dyads (13 male 31 female) who mutually reported not clicking (19 dyads) or had one-sided click (25 dyads). A leave-one-out cross-validation gave rise to a meaningful receiver operator curve (ROC) (Fig. 4). The area under the curve (AUC) was 0.67, which is significantly different from chance (U = 2.811, p = 0.0049). This reflects a cross-validation accuracy of 71.21% (binomial p = 0.00076, Cohen’s g = 21.21%), permitting correct identification in 17 of 22 mutual click reports and 30 of 44 reports of no mutual click (77.27% sensitivity and 68.18% specificity).

**Fig. 4.**
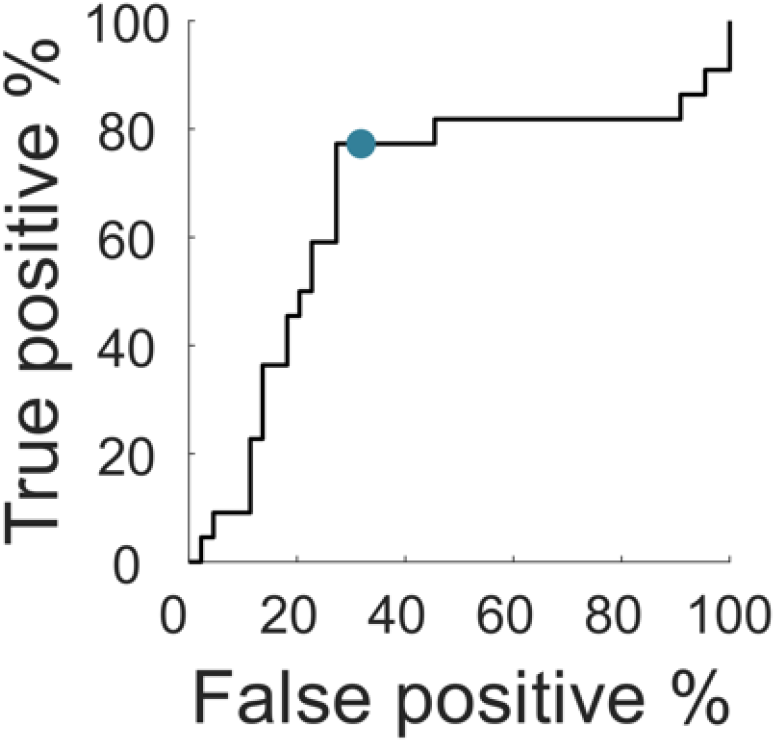
Classifying mutual click dyads by eNose-derived body-odor similarity. An ROC classifying dyads who mutually clicked (n = 22, 14 females) or didn’t mutually click (n = 44, 31 females). The blue dot marks the performance of the leave-one-out cross-validation weighted KNN classifier.

## Discussion

> We sometimes encounter people, even perfect strangers, who begin to interest us at first sight, somehow suddenly, all at once, before a word has been spoken.
>
> — Fyodor Dostoevsky, Crime and Punishment, 1866

Here we investigated an alternative hypothesis, namely that perfect strangers may begin to interest us at first sniff, rather than first sight alone. Across experiments, our data converged to imply that the body-odors of same-sex click-friends are more similar to each other than the body-odors of same-sex random dyads. Olfaction is a dominant sensory input underlying social interaction (*19*), and this statement is rather unarguable for all terrestrial mammals but one: Humans. In humans, the role of olfaction has been denigrated primarily because of various social taboos (*37*), culminating in the view that olfaction is unimportant for human sociality (*38*). Recent evidence, however, implies a significant role for olfaction in human social interaction, albeit a role that materializes mostly without conscious awareness. Humans are constantly but mostly subconsciously sniffing themselves (*21*) and their conspecifics (*20*). These odors then have a host of effects, and may carry a host of information. We will note a few standout examples beyond those detailed in the introduction: Sniffing women’s body-odors coordinates women’s menstrual cycles (*39*) (although this effect remains debated (*40*)). Sniffing women’s tears lowers testosterone in men (*41*, *42*), which inevitably alters behavior. Sniffing one particular molecule expressed in body-odor (Androstadienone) raises levels of cortisol in women (*43*), and sniffing a different particular molecule expressed in body-odor (Hexadecanal), blocks aggression in men, but triggers aggression in women (*44*). Humans can infer a state of disease in body-odor (*45*), and can smell aggression (*46*), fear (*25*), stress (*47*), depression (*48*) and happiness (*26*). Body-odor also serves as a cue for human kinship (*22*, *24*) and influences human mate choice (*49*). In the current study we add to this by finding that humans may use olfactory information to guide preferences in same-sex non-romantic dyadic interactions. This notion is consistent with reports on impaired sociality in human congenital anosmia (*50*), and on altered social chemosignaling in autism spectrum disorder (*51*).

This study has several limitations we would like to acknowledge. First, we placed our research in the context of click-friendships, but did not enter other types of friendship as a control. In other words, we find that body-odor similarity is greater between click-friends versus random dyads, but we did not test this versus “ordinary” friend dyads that were not click-friends. Thus, body-odor similarity may be a component of all friendships, not click-friendships alone. Second, we acknowledge the two null results obtained in this study: In the first, perceived body-odor similarity was not different across click and random dyads in the triangle test of Experiment 3. We think that the odor-memory-related difficulty of this task underlies this null result, and we can add that visual similarity, despite it being a well-known friendship similarity cue (*9*), was also not related to friendship when using the triangle paradigm. Despite this mitigating factor, this null result implies limited effect magnitude. The second null result was in the motion energy analysis of the mirror game in Experiment 5. This would have provided a valuable objective measure in this task, and in its absence, we rely on subjective measures alone to analyze the mirror-game results. This in contrast to Experiment 3 for example, where sniff patterns provided for an objective similarity measure to bolster subjective perception. Finally, although not a limitation per se, we would like to emphasize that the chemical similarity that was inferred by eNose was unrelated to the perceptual similarity inferred by human raters. In other words, these measures captured independent sources of variance, and this keeps us further away from identifying what components of body-odor ultimately contribute to the observed effects. Indeed, although the eNose is a convenient tool, its information is limited, and hard to generalize. For example, although eNose similarity was related to the quality of dyadic interaction in two separate experiments, we observe that the absolute eNose values were very different across these two experiments that were conducted more than a year apart. Thus, the eNose can report on chemical similarity within an experiment, but remains largely uninformative beyond this.

And yet, despite these limitations, this study also has specific strengths, and primarily the recurrence of the link between body-odor similarity and quality of same-sex dyadic interaction across multiple experiments and experimental designs. Moreover, we think that the predictive eNose model applied to the round-robin interaction between strangers in Experiment 5 demonstrates causality, that is, not only do friends smell more similar to each other than expected by chance, but also that humans use this information to make decisions on strangers they meet for the very first time. This result may have profound technological implications, for example, use of an eNose to pair individuals for a dyadic task. Here, however, we prefer to ask what is the weight of this in real life? We do not know. Our round-robin experiment diverged from real life in that participants were not allowed to speak with each other. In the real world, humans use complex language to interact, and it is in this that we are indeed most different from other terrestrial mammals in our social interactions. Nevertheless, we think our results imply that we may also be more like other terrestrial mammals in this respect than we typically appreciate. This appreciation is important, because beyond a deeper understanding of human behavior, it may point towards novel paths to intervention in social impairment. We conclude in reiterating that several experiments converged to suggest that human same-sex non-romantic friends smell more similar to each other than expected by chance, and that complete strangers who smell more similar to each other as determined by an eNose then have better dyadic interactions. Thus, humans may use their nose to sniff out new friends.

## Acknowledgment

This work was funded by an ERC AdG grant (SocioSmell 670798) awarded to Noam Sobel. Inbal Ravreby is funded by the Ariane de Rothschild Women’s Doctoral Program. We thank L. Rozenkrantz for insightful discussion.

## Author Contributions

Conceived the idea: IR. Designed experiments: IR; NS. Ran experiments: IR. Analyzed data: IR; KS. Wrote first draft: IR. Edited final draft: IR; KS; NS.

## Supplementary Materials

Materials and Methods

Table S1

Fig S1 – S4

## Materials and Methods

### Participants

All participants provided written informed consent to procedures approved by the Wolfson hospital Helsinki Committee (protocol reference number 0035-16-WOMC). All participants were screened for self-reported lack of nasal congestion or olfactory dysfunction, and then participated for monetary reward. Distribution of participants across experiments was as follows: **Define “click-friendship”:** We recruited 235 participants online, 135 females and 100 males aged between 20 and 42 years (M = 26.35 ± 4.166). **Harvesting click dyads’ body-odor:** To find click-friends, we posted extensively on campus billboards and on social media. Respondents were first phone-interviewed, and then interviewed by questionnaire to verify that they satisfied click-friend criteria. This recruitment effort lasted 6 months, and entailed collecting body-odors from participants across the entire country. In total, we harvested body-odor from 20 pairs of same-sex close friends (half males and half females) who reported that their friendship began as a “click-friendships”, aged between 22 and 39 years (M = 24.757 ± 3.388, mean friendship duration = 6.185 ± 5.793 years). **Triangle test and ratings:** We recruited 24 naive participants, 13 females, aged between 22 and 39 years (M = 27 ± 4.63). **Explicit similarity ratings:** We recruited for olfactory similarity ratings 25 participants, 19 females, aged between 21 to 38 years (M = 25.76 ± 4.075). For visual similarity ratings we recruited another 25 participants, 13 females, aged between 21 to 37 years (M = 26.56 ± 5.091). **The Mirror Game:** We recruited a total of 17 naive healthy participants that did not know each other personally, 10 females, aged between 20 and 37 years (M = 26.2 ± 4.56). No prior acquaintance between dyads was verified through detailed questionnaires where it was uncovered that 2 of the 21 male dyads and 5 of the 45 female dyads had attended common undergraduate large-scale courses, but they had no personal interaction, and proclaimed not to know each other.

### Paradigms

#### Define “click-friendship”

The participants were asked to define “click-friendship” in their own words (in Hebrew). Ten participants responded that they cannot, retaining 225 respondents.

### Harvesting body-odor from click-dyads

Donors were provided with non-perfumed soap to shower with each evening before wearing provided 100% cotton T-shirts to wear on two consecutive nights. Donors were instructed to use only the items provided to them, and avoid other soaps and the use of lotions, deodorants, antiperspirants, perfumes, colognes, etc. They were instructed to wear the shirts at least six hours each night, and prevent other humans or pets from sleeping on, or using, the bed during the testing period. Moreover, they were asked to avoid foods that strongly influence body-odor such as curry, amchoor, fenugreek, asparagus and garlic. In the morning after the first night, donors placed their shirts in a plastic zip-lock bag to prevent absorption of other odors and to keep the participants’ body-odor in the shirts. After the second night the donors were instructed to store the bagged shirts in the freezer to minimize loss of odor. The T-shirts were collected (typically on the same day) and stored in lab at −20°C in designated glass jars. Each donor also completed a general questionnaire.

#### eNose similarity test

To examine whether there is chemical similarity between click dyads’ body-odor, we used a PEN3 eNose (AIRSENSE Analytics GmbH, Schwerin, Germany). The PEN3 is a compact (92 × 190 × 255 mm) lightweight (2.3 kg) device, consisting of a gas sampling unit and a sensor array. The sensor array is composed of 10 different thermo-regulated metal oxide sensors, positioned in a stainless-steel chamber (volume: 1.8 ml, temperature: 110 °C). Each sensor is uniquely coated, rendering it particularly sensitive to a restricted class of chemical compounds. When a compound interacts with the sensor, this results in an oxygen exchange that leads to a change in electrical conductivity (*52*). We used the PEN3 with its native sampling software (WinMuster), and the following settings: Chamber flow = 400ml/min, Flush time = 100s, Zero-point trim time = 10s, Measurement time = 80s. T-shirts were first thawed for 1 hour in room temperature. Then, in order to measure the headspace, we covered each jar with parafilm sheet and waited another 1 hour. Next, we used the eNose to measure the headspace of each jar at room temperature (see Fig. 1A).

#### Triangle test

The 40 T-shirts of the click dyads were thawed at room temperature 1 hour before the smelling experiment. We used a previously described shirt sniffing device (SSD) (*53*) to standardize body-odor sampling. The SSD consists of a glass jar containing the T-shirt, with an air intake port via soda lime filter, and air sampling port via one-way flap valve into individual-use airtight nose mask (*53*). Using the SSD assured that environmental odors, and/or other participant odors, did not contaminate the sample. To probe explicit classification, we used a triangle test. On each trial, the participants were presented with a body-odor triplet: two odorants were from a click dyad, and the third distractor odorant was from an unrelated same-sex body-odor donor. Participants were asked to select the odorant outlier. Each participant completed 20 trials (inter-trial-interval = 25 seconds), one for each click dyad. The triangle smell test was followed by a control triangle visual test, in which participants were asked to select the outlier picture, according to pictures of the same three people, randomly ordered. To match the odor test, in which the odors in each triplet were smelled one after another, the pictures were presented one-by-one rather than simultaneously side-by-side. During sampling, we measured nasal airflow using a nasal cannula (1103, Teleflex medical) placed at the nares and attached to a spirometer (Spirometer FE141 ADInstruments). The nasal airflow was sampled at 1 kHz and recorded using a Power-Lab 16SP Monitoring System (ADInstruments, Australia). Airflow data were later displayed, stored, reduced and analyzed using LabChart 7 software (ADInstruments).

#### Descriptor ratings

Each participant smelled the 40 body-odors one-by-one, randomly ordered, and rated them using visual analog scales (VASs) for “pleasantness”, “intensity”, “sexual attraction”, “competence” and “warmth” (temperament). Nasal airflow was monitored throughout this task as before.

#### Explicit similarity ratings

Each participant was presented with 40 dyads, 20 consisting click-friends and 20 random dyads. They rated the odors of each dyad using a VAS ranging from “different” to “similar”.

#### The Mirror Game

The participants were asked to avoid use of perfume or deodorant and to avoid foods known to influence body-odor (as detailed previously). The experiment took place in two different round robin sessions, one for females and one for males. In each session the participants were requested to split into pre-assigned dyads in each round. All pairs were tested simultaneously in separate identically arranged experimental rooms. In each round the dyads were instructed to stand facing each other, at a distance of 50 cm, which was marked on the floor. This minimal distance ensured exposure to conspecific volatiles. Then the dyads were asked to play the full body Mirror Game, in which they had to move their hands coordinately while keeping their legs at the starting point, with no designated leader or follower (Fig. 2A). The participants were not allowed to speak with each other during the entire experiment. In this way, the impression formation was not influenced by voice or by a conversation content, but only by the non-verbal interaction. During the game the participants were filmed using a hidden camera. Each game round lasted 2 minutes and each participant played the Mirror Game with all the other same sex participants, culminating in 45 female dyads and 21 male dyads. After each round the participants were asked to use an on-screen indicator where they freely moved circles that denoted themselves and their partners towards or away from each other, according to their feelings of “closeness” to and “overlapping” with their partner in the Mirror Game. This was used to estimate closeness in terms of self-other boundaries, an adaptation of the Other in the Self (IOS) scale (*36*). The distance between the other and the self was calculated by extracting the distance (in pixels) between the centers of the two circles. In addition to the IOS scale, the participants were requested to indicate whether they had a “click” with their partner or not. Additionally, each participant was asked to indicate on a VAS of 1-100 the following aspects regarding clicking: how much they read their partner’s mind, understood their partner, would like to meet again with their partner, wanted to know their partner, felt close to their partner, liked their partner, felt chemistry with their partner, thought that they could be good friends, felt on the same wavelength, had a feeling that they already knew their partner, felt comfortable to share personal issues with their partner, felt that their partner was friendly toward them.

#### Click-dyad classification by eNose

We used a leave-one-out weighted KNN classifier in order to classify dyads who reported mutual click vs. dyads who did not report mutual click. The classifier was applied to the difference between the five activated sensors (# 2, 6, 7, 8, 10) of each member in a dyad. To have a consistent geometry, we kept the first component of the vector positive (i.e. multiple the vector by minus one if the first component was negative). We then used PCA, keeping 95% of the variance, and thus reduced the five dimensions into two. We then classified the resulting data with the following parameters: the number of neighbors was 13, the distance metric was Euclidean distance and the distance weight was squared inverse. To correct for the imbalanced dataset (22 dyads that mutually clicked vs 44 dyads that did not mutually click), we used cost-sensitive learning during the training. The cost was equal to the inverse of the proportion between mutual-click dyads and no-mutual-click dyads.

### Statistics and inclusion/exclusion criteria

#### Data analyses software

All data analyses were performed using MATLAB R2018a and JASP (Version 0.13.1.0).

#### Define “click-friendship”

**Statistics:** The frequencies of the statements. **Exclusion:** Ten participants out of the 235 participants (4.26%) did not know what “click-friendship” means and thus were excluded.

#### Triangle test

**Statistics:** One sample two tailed t-test of the d-prime (d’) scores against d’ = 1. **Exclusion:** There were six trials of missing data (out of 480 trials overall, i.e., 1.25%), in which the participants mistakenly inserted odor numbers that did not exist.

#### Descriptors ratings

**Statistics:** To account for individual differences in use of scales and for differences in use of different descriptors, each participant’s data were normalized per descriptor by first subtracting the minimal value applied by the participant, and then dividing by the maximal remaining value. This generated a normalized range for each descriptor, between 0 and 1. To obtain a five-dimensional VASs ratings’ vector we used the average rating in each descriptor for each body-odor of the 20 click dyads, and then calculated the Euclidean distances. We used the 40 individuals in the click dyads cohort to randomly generate 10,000 iterations of 20 non-click-same-sex dyads. In each iteration we averaged the Euclidean distances of the 20 random dyads, to obtain a distribution of the mean Euclidean distances between 20 random dyads, 10,000 times. We then evaluated where the mean Euclidean distance of the 20 click dyads falls in the distribution. **Exclusion:** One participant out of the 24 dropped out mid-experiment. In three participants movement of the nasal cannula prevented nasal airflow analysis.

#### Explicit similarity ratings

**Statistics:** The ratings between “different” and “similar” were first transformed to a scale between 0 and 100. Then, to account for individual differences in use of scales, we normalized the data and compared between the click dyads and random dyads the same way as we did for the five descriptor ratings. **Exclusion:** there were no exclusions.

#### The Mirror Game

**Statistics:** To test whether the 22 dyads that mutually reported clicking with each other have a body-odor that is chemically more similar than the body-odor of the other 44 dyads, we compared the eNose distance between these 22 dyads to the eNose distance between 10,000 random selections of 22 (out of the 66) same-sex non-mutual-clicking dyads. To get dyadic reports out of the 13 self-reports, we averaged each self-report of the two partners in a same-sex dyad. We used a Pearson correlation to examine the relationship between various aspects of social interaction quality and the chemical body-odor similarity. We asked a naive judge to watch the videos of the mirror game one by one, to verify that indeed the participants played as if they tried to mirror each other. **Exclusion:** In the correlation analyses, the exclusion criteria threshold was 2.5 SD from the regression line. Accordingly, out of the 66 dyads, two dyads (3%) were excluded from the IOS, one from reading the partner’s mind, one from understanding the partner, one from willingness to meet again, one from willingness to know the partner, one from feeling close to the partner, two from liking the partner, one from feeling that there in chemistry with the partner, one from thinking that they could be good friends and one from feeling that the partner was friendly.

#### eNose classification of Click dyads

**Statistics:** We used a Mann–Whitney U test to examine whether the area under the ROC is significantly different from chance. A binomial test was used to examine whether the classification accuracy was different from chance. To estimate the success of the classifier we also calculated sensitivity and specificity. **Exclusion:** there were no exclusions.

