## Supplemental Data 1 for "Sniffing Out New Friends: Similarity in Body-Odor Predicts the Quality of Same-Sex Non-Romantic Dyadic Interactions"

### Supplementary Materials

**Table S1. Statements frequencies in "click friendship" definitions defined by 225 participants.**

| Frequency | Statement |
|---|---|
| 137 | Immediately when meeting / from first glance / right away |
| 40 | Matching |
| 26 | Good friendship that is rapidly formed |
| 23 | Chemistry |
| 22 | Understanding each other |
| 22 | "On the same wavelength" |
| 17 | Mutual |
| 17 | A feeling of knowing each other for a long time from the first interaction |
| 17 | Flowing conversations |
| 17 | Common language / common ground |
| 16 | Strong friendship |
| 15 | Shared point of view / opinions |
| 14 | Common interests |
| 14 | Effortless interaction |
| 12 | In the first meeting it is possible to forecast that we will be friends |
| 12 | Comfort with each other from the beginning / openness / feeling free to talk about anything |
| 9 | Many topics for conversation |
| 8 | Non-verbal understanding / no need to explain oneself |
| 8 | Unexplained bonding / magical |
| 7 | Closeness |
| 7 | Bonding |
| 6 | Curiosity to know the other, willingness to deepen the interaction and become friends |
| 6 | Joy |
| 5 | Love |
| 5 | Physical or emotional attraction |
| 4 | Making each other laugh |
| 4 | Deep / profound |

| | |
|---|---|
| 4 | Willingness to be together |
| 4 | Talking a lot, spending a lot of time together |
| 4 | A sparkle in the eyes, fire in the eyes |
| 4 | Honesty |
| 3 | Rare |
| 3 | Trying to impress |
| 2 | Excitement |
| 2 | Might not be real / not last long |
| 2 | Finding a pattern we look for |
| 1 | Falling in love |
| 1 | Acceptance |
| 1 | Sharing without judging |
| 1 | Identifying the advantages of the other |
| 1 | Empathy |
| 1 | When the eyes meet |

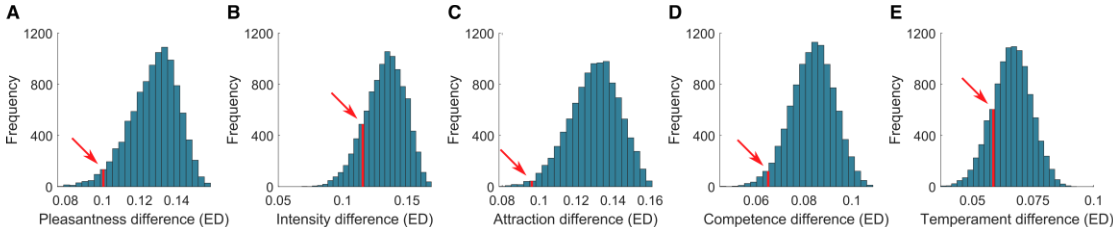


**Fig. S1. Histograms showing 10,000 iterations of the average Euclidian distance between 20 same-sex random dyads, as represented by VAS ratings in five descriptors.**

The average Euclidian distance for the 20 click dyads is indicated in each histogram by the red arrow pointing on the red line. In each histogram half of the random dyads are males and half are females. (**A**) click-dyads were rated as significantly more similar in body-odor pleasantness than the odor of random dyads (mean clicking = 0.107 ± 0.12 11 AU, mean random = 0.135 128 ± 0.013 0.014 AU, bootstrapped p = 0.03703, Cohen's d = 2.1542). (**B**) There was no significant difference in the intensity ratings similarity between click dyads and random dyads (mean clicking = 0.116 ± 0.095 AU, mean random = 0.133 ± 0.0154 AU, bootstrapped p = 0.143, Cohen's d = 1.104). (**C**) Click dyads' body-odor was rated as significantly more similar in attractiveness than the body-odor of random dyads (mean clicking = 0.1 096 ± 0.114 106 AU, mean random = 0.136 ± 0.014 013 AU, bootstrapped p = 0.00901, Cohen's d = 2.6152.571). (**D**) Click dyads' body-odor was rated as significantly more similar in its association to competence than the body-odor of random dyads (mean clicking = 0.065 ± 0.063 AU, mean random = 0.084 ± 0.009 AU, bootstrapped p = 0.027, Cohen's d = 2.111). (**E**) Click dyads' body-odor was not rated as significantly more similar in its association to warmth than the body-odor of random dyads (temperament) (mean clicking = 0.059 ± 0.041 AU, mean random = 0.066 ± 0.008 AU, bootstrapped p = 0.175, Cohen's d = 0.875).

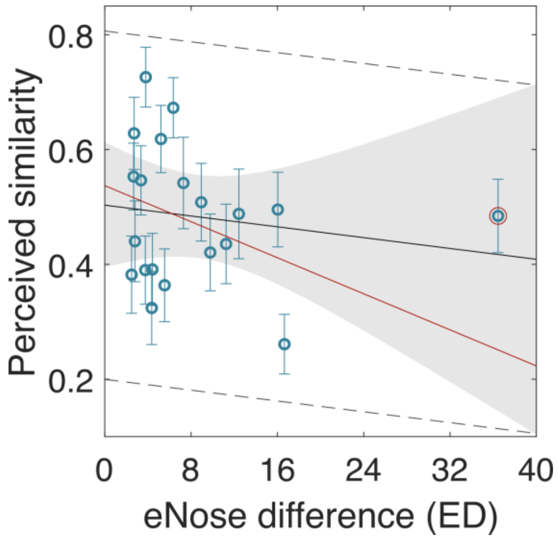


**Fig. S2. Pearson correlation between dyad similarity ratings as obtained with eNose and perception**

Pearson correlation showed no significant relationship (r = -0.16, p = 0.51, and after removing the right outlier which is circled in red, r = -0.29, p = 0.23). The blue circles represent each of the 20 click-dyads. The black line is the linear regression line and the grey area marks the CI of the regression line. The dashed black lines denote a distance of ± 2.5 SD from the regression line. The red line is the linear regression line without the outlier.

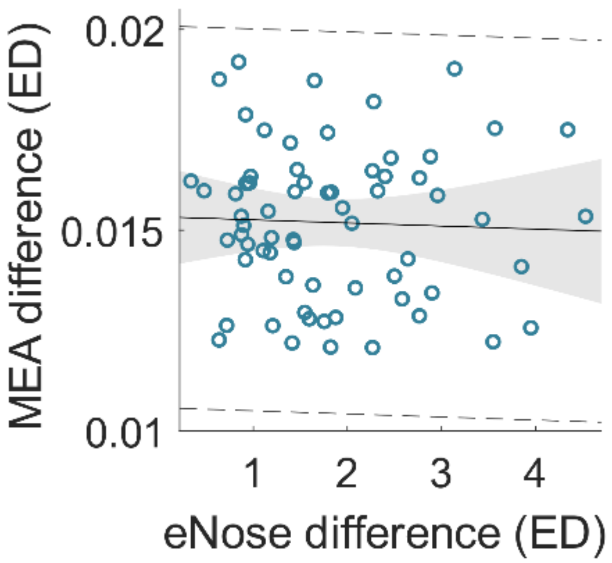


**Fig. S3. Pearson correlation between synchronization level in the mirror game (motion energy difference) and body-odor chemical similarity (eNose difference).**

Pearson correlation showed no significant relationship ($r = -0.04$, $p = 0.75$). The blue circles represent each dyad that played the mirror game. The black line is the linear regression line and the grey area marks the CI of the regression line. The dashed black lines denote a distance of ± 2.5 SD from the regression line. A naive judge that was instructed to watch the videos one by one concluded after watching all the 66 videos that it seems that the females highly tried to be in sync whereas many and maybe even most of the males mainly tried to challenge each other, more than they tried to keep being in sync (notice that we did not ask the judge to judge males' and females' games separately, this was a spontaneous observation). This difference between the way males and females played the mirror game may be the reason that there was no correlation between the synchronization level and the body-odor chemical similarity.

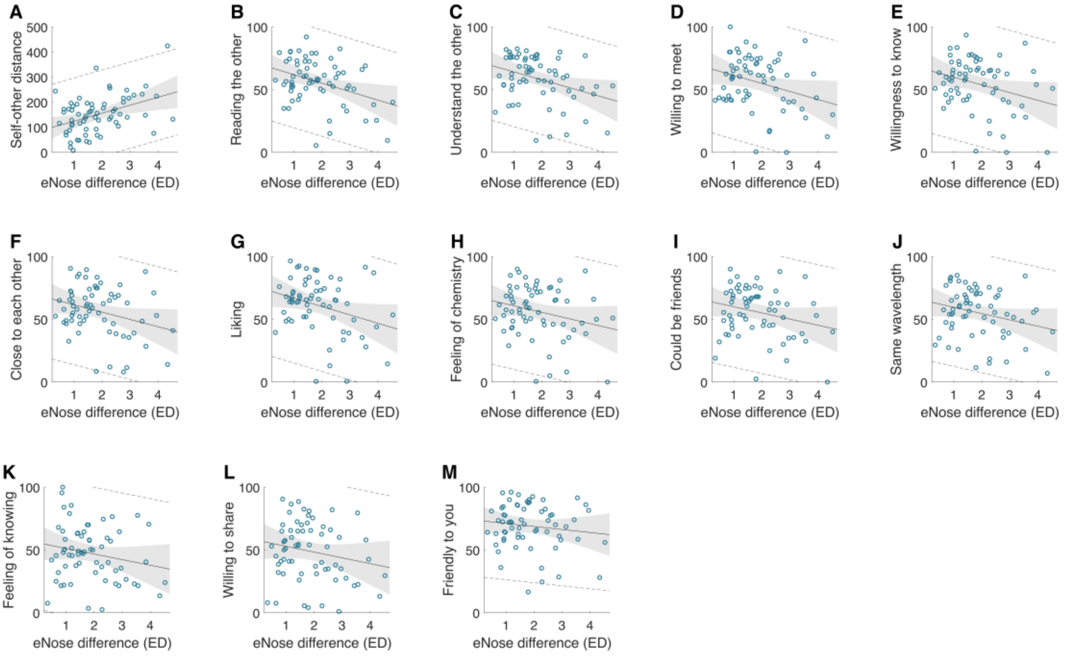


**Fig. S4. Predicting various aspects of social interaction quality by body odor similarity (eNose difference) without excluding outliers.**

The blue circles represent each dyad that played the mirror game. The black line is the linear regression line and the grey area marks the CI of the regression line. The dashed black lines denote a distance of ± 2.5 SD from the regression line – the exclusion threshold criteria. The figure show correlations between eNose ED and (A) including the other in the self as was measured in the IOS scale by pixels distance ($r = 0.419$, $p = 0.0005$), (B) reading the partner's mind ($r = -0.37$, $p = 0.0021$), (C) understanding the partner ($r = -0.342$, $p = 0.005$, (D) willingness to meet again with the partner ($r = -0.3$, $p = 0.0145$), (E) willingness to get to know the partner ($r = -0.296$, $p = 0.016$), (F) feeling close to the partner ($r = -0.292$, $p = 0.017$), (G) liking the partner ($r = -0.308$, $p = 0.012$), (H) feeling chemistry with the partner ($r = -0.249$, $p = 0.044$), (I) thinking that they could be good friends ($r = -0.246$, $p = 0.046$), (J) feeling on the same wavelength ($r = -0.26$, $p = 0.036$), (K) feeling as if they already knew the partner ($r = -0.21$, $p = 0.098$), (L) feeling comfortable to share personal issues with the partner ($r = -0.2$, $p = 0.11$) and (M) feeling that the partner was friendly ($r = -0.133$, $p = 0.29$).
